# N-Amino Peptide-Graphene Quantum Dot Loaded Small Extracellular Vesicles for Targeted Therapy of Tauopathies

**DOI:** 10.1101/2024.12.23.630154

**Authors:** Runyao Zhu, Gaeun Kim, Benjamin H. Rajewski, Isaac J. Angera, Juan R. Del Valle, Yichun Wang

## Abstract

Tauopathies, a group of neurodegenerative disorders, are characterized by the abnormal aggregation of tau proteins into neurofibrillary tangles (NFTs), driving synaptic dysfunction, neuronal loss, and disease progression through tau aggregate propagation. Graphene quantum dots (GQDs) functionalized with *D*- cysteine (*D*-GQDs) have shown promise in inhibiting tau aggregation and transmission *via* π-π stacking and electrostatic interactions with tau proteins. However, the non-specific binding of GQDs to various proteins in the physiological environment, such as serum albumin, limits their clinical translation. In this study, we aim to enhance the specificity of *D*-GQDs toward tau protein by incorporating a tau-targeting N-amino peptide, mxyl-NAP2. The mxyl-NAP2/*D*-GQD complex demonstrated improved selectivity for tau protein over serum albumin, effectively enhancing the inhibition of tau aggregation. To further minimize off-target effects and optimize therapeutic delivery, we loaded the mxyl-NAP2/*D*-GQD complex into small extracellular vesicles (sEVs), followed by functionalization of sEVs with neuron targeting ligand, rabies viral glycoprotein peptides. This strategy not only reduced off-target effects, but also enhanced uptake by neuron cells, which further improved inhibition of tau transmission between neurons. Our results indicated that mxyl-NAP2/*D*-GQD-loaded sEVs hold great promise for overcoming the off-target limitations of *D*- GQDs and advancing the development of precision therapeutics for neurodegenerative diseases.

## 1. Introduction

The pathological aggregation of the microtubule-associated protein tau into neurofibrillary tangles (NFTs) is a hallmark of various neurodegenerative diseases, collectively known as tauopathies, including Alzheimer’s disease (AD),^1^ corticobasal degeneration (CBD),^2^ and progressive supranuclear palsy (PSP).^3^ Tau is an intrinsically disordered protein that plays a critical structural and regulatory role in neurons, stabilizing microtubules, maintaining neuronal cytoskeleton, and modulating cellular signaling.^4, 5^ However, under pathological conditions, abnormal post-translational modifications and certain environmental stimuli can lead to the dissociation of tau from microtubules and its aggregation into insoluble tau filaments. These aggregates, deposited in the brain, are strongly associated with synaptic dysfunction, neuronal loss, and brain atrophy.^4, 6, 7^ In addition, tau oligomers and fibrils can propagate from diseased to healthy neurons, inducing further tau aggregation in a prion-like manner and leading to the spread of pathology.^8, 9^ Over the past decades, various therapeutic strategies have emerged based on the pathogenesis of tauopathy, such as small molecules designed to modulate tau post-translational modifications,^10, 11^ inhibit tau aggregation,^12, 13^ or promote fibril degradation,^14, 15^ as well as anti-tau antibodies for active and passive immunotherapy,^16, 17^ aimed at blocking the spread of tau aggregates and promoting their clearance from the brain.

Nanoparticles (NPs), including polymeric NPs, lipid NPs, and carbon NPs, have been investigated as a therapeutic strategy for treating neurodegenerative diseases.^18–20^ Graphene quantum dots (GQDs), as a subclass of carbon NPs, exhibit lateral dimensions typically smaller than 20 nm and are composed of one or a few atomic layers of graphene.^21, 22^ Their unique properties, including tunable photoluminescence, excellent photostability, small size, high biocompatibility, low cytotoxicity, and rapid renal clearance, make them ideal candidates in advanced biomedical applications, such as bioimaging, photodynamic therapy, and drug delivery.^23–25^ Beyond these general attributes, GQDs possess a distinctive π-conjugated planar structure that allows them to interact with aromatic amino acids in tau proteins *via* π-π stacking, including histidine, phenylalanine, and tyrosine.^26, 27^ Building on this characteristic, we previously developed the functionalized GQDs with *D*-cysteine (*D*-GQD) for the treatment of tauopathy.^28^ *D*-GQDs exhibit negative surface charge, which further facilitates electrostatic interactions with positively charged aggregation-prone domains of tau. These synergistic interactions enable *D*-GQDs to disrupt pathological tau processes— hindering its aggregation, promoting tau fibril disassembly, and preventing cell-to-cell propagation of tau aggregates. Despite their promising potential, these GQDs face challenges in translational applications. Their biomimetic surface chemistry, while advantageous in certain interactions, may result in non-specific binding to other proteins, such as other amyloids,^29–31^ and human serum albumin (HSA),^32^ which potentially reduce their specificity and efficacy *in vivo*.

To address the challenge of the non-specific binding of GQDs to proteins in the physiological environment, conjugating GQDs with bioactive ligands specifically targeting tau protein—*via* either covalent or noncovalent bonds—offers a powerful way to enhance molecular recognition and improve therapeutic efficacy.^33, 34^ In our previous work, we developed a class of macrocycles, termed β-bracelets, which mimic the cross-β epitopes common to pathological tau filaments. ^35–37^ Backbone N-amination of the aggregation-prone hexapeptide module (PHF6; _306_VQIVYK_311_) within these β-bracelets affords soluble inhibitors of aggregation and propagation. Herein, we combine a tau-targeting β-bracelet ligand, mxyl-NAP2, with the highly efficient inhibitory effect of GQDs, providing a more effective solution for selective inhibition of tau aggregation.

Another effective strategy to reduce the off-target effects of GQDs involves encapsulating them within targeted drug delivery carriers, which enhances specificity and minimizes toxicity to non-target tissues.^38,39^ Among nanocarriers, lipid-based nanoparticles have been extensively investigated for drug delivery applications due to their biocompatibility and ability to encapsulate both hydrophobic and hydrophilic therapeutics.^40^ Small extracellular vesicles (sEVs), lipid NPs naturally secreted by most eukaryotic cells, have emerged as a promising alternative for drug delivery.^41^ These sEVs, approximately 50∼150 nm in size, can naturally transport bioactive molecules, such as proteins and nucleic acids, to recipient cells, making them major mediators of intercellular communication.^42^ Furthermore, the surfaces of sEVs can be further functionalized with specific ligands using chemical or biological engineering techniques, thereby improving their precision in targeting diseased tissues.^43–45^ Most importantly, our recent findings reveal that chiral GQDs, especially *D*-GQDs, exhibit high permeability into sEV. This unique property is attributed to favorable chiral interactions with biological lipid membranes, allowing efficient encapsulation of GQDs within the vesicles without alternating lipid membranes of sEVs.^23, 46, 47^ By combining the encapsulation and targeting capabilities of sEVs with the functional properties of chiral GQDs, this hybrid approach offers a highly adaptable platform for therapeutic delivery, reducing off-target effects while enhancing treatment efficacy for tauopathies.

In this study, we developed mxyl-NAP2/*D*-GQD-loaded sEVs as a targeted delivery system for the effective treatment of tauopathies **(Fig. 1)**. To improve the selectivity of *D*-GQDs for tau protein, we first incorporated *D*-GQDs with mxyl-NAP2, a type of N-amino peptide (NAP) β-bracelet that targets the aggregation-prone PHF6 motif in tau protein, forming a mxyl-NAP2/*D*-GQD complex.^36^ By monitoring tau aggregation in the presence of HSA, we found that this complex significantly enhanced the selectivity of *D*-GQDs for tau over HSA while effectively inhibiting tau aggregation. Furthermore, we encapsulated the mxyl-NAP2/*D*-GQD complex into sEVs, leveraging the superior permeation efficiency of *D*-GQDs into sEVs. A cellular propagation assay employed to test the tau propagation demonstrated that the mxyl-NAP2/*D*-GQD-loaded sEVs achieved more efficient inhibition of tau fibril propagation. Additionally, we engineered the sEV surface with rabies viral glycoprotein (RVG) peptides, a targeting ligand to neurons, facilitating targeted delivery to neurons and further reducing tau transmission between cells. Overall, our findings suggest that the mxyl-NAP2/*D*-GQD complex, especially when targeted delivered *via* sEVs to neurons, offers a promising targeted and effective therapeutic strategy for tauopathies.

**Fig. 1.**
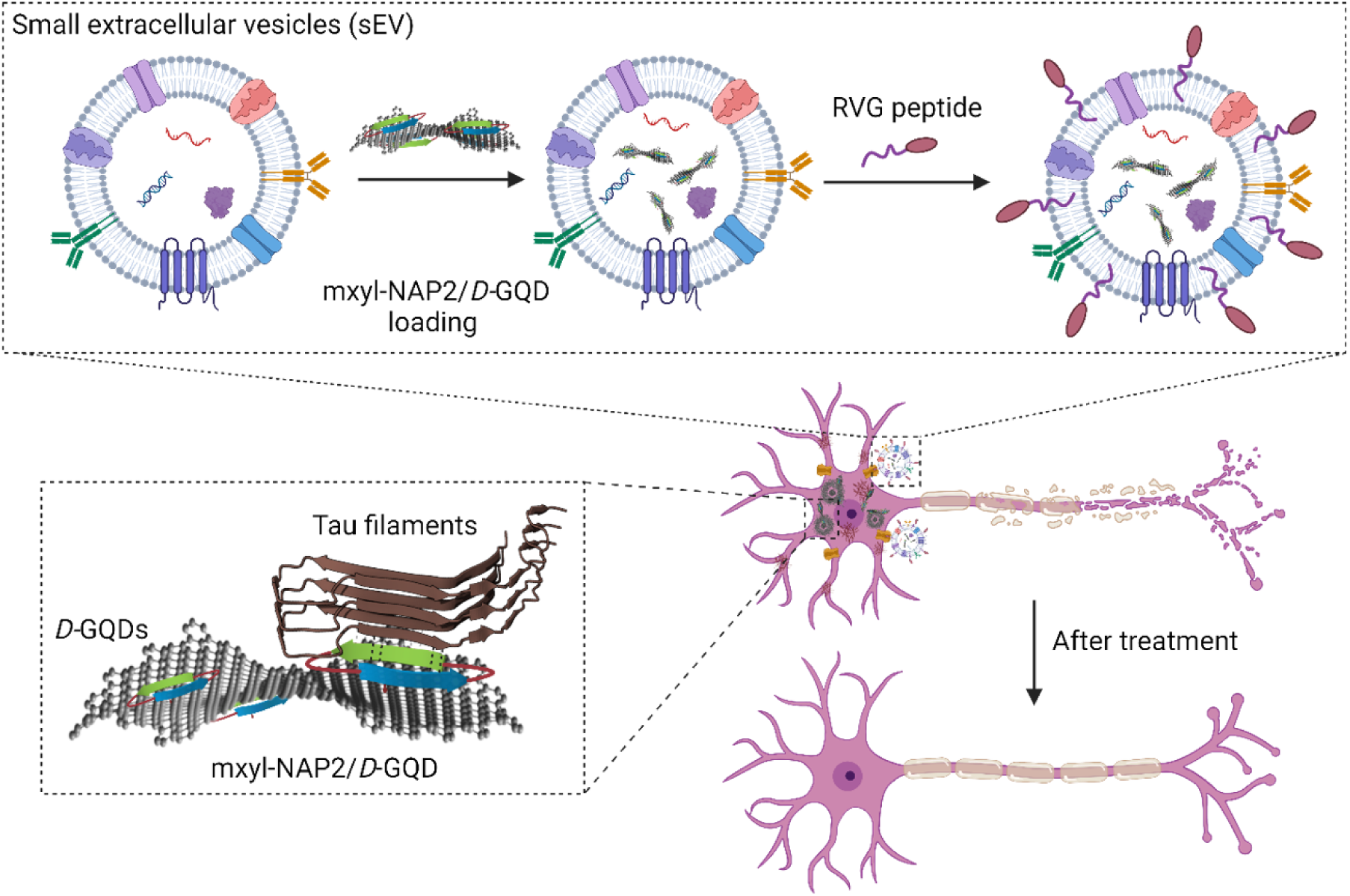
Schematic illustration of RVG-mxyl-NAP2/*D-*GQD-sEV with the ability to inhibit the seeding activity of tau fibrils for tauopathy therapy. *D*-cysteine functionalized GQDs (*D*-GQD) incorporate mxyl-NAP2 that binds specifically with PHF6 hexapeptide in tau filaments (PDB 6QJH), enhancing the selectivity of *D*-GQDs. Rabies viral glycoprotein (RVG) peptide-modified small extracellular vesicles (sEVs) can be recognized by nicotinic acetylcholine receptors (nAchR) on neurons, increasing their cellular uptake.

## 2. Results and discussion

### 2.1. Incorporation of *D*-GQD with tau-targeting NAP *via* π-π stacking

As-synthesized GQDs carrying carboxyl groups on the edges were successfully obtained following the previously reported method **(Fig. 2a)**.^48^ We then functionalized GQDs with *D*-cysteine, which resulted in *D*-GQDs with a significant inhibition effect on tau aggregation^28^ and high permeation efficiency into sEVs reported in our previous works **(Fig. 2b)**.^23, 46^ The structures of GQDs and *D*-GQDs were confirmed by the combination of spectroscopy and microscopy. The average diameter of *D*-GQDs was 7.99 ± 2.03 nm, detected by transmission electron microscopy (TEM) **(Fig. 2c-d)**. In the circular dichroism (CD) spectra **(Fig. 2e)**, *D*-GQDs showed a negative peak at 213 nm, close to the chirality peak of free *D*-cysteine at 209 nm. Additionally, *D*-GQDs gave rise to a cloud peak with an opposite sign at 258 nm indicating covalent bonding of *D*-cysteine on the edge of GQDs, whereas pristine GQDs displayed no chiroptical activity in CD spectra.^47^ In the fourier transform infrared (FTIR) spectra **(Fig. 2f)**, due to the carboxyl groups in the structure of *D*-cysteine, the peak at 1706 cm^-1^ assigned to C=O stretching was more obvious in the spectrum of *D*-GQDs than that of GQDs.^48^ There was also a peak that appeared at 1250 cm^-1^ in *D-*GQD samples, attributed to C-N stretching.^49^ The zeta potential of *D*-GQDs was changed to-1.46 ± 0.42 mV from-19.9 ± 2.30 mV of GQDs. Due to the electron-donor, *D*-cysteine, the fluorescence emission spectrum of *D*-GQDs was red-shifted, compared to GQDs **(Fig. S1)**.^50^

**Fig. 2.**
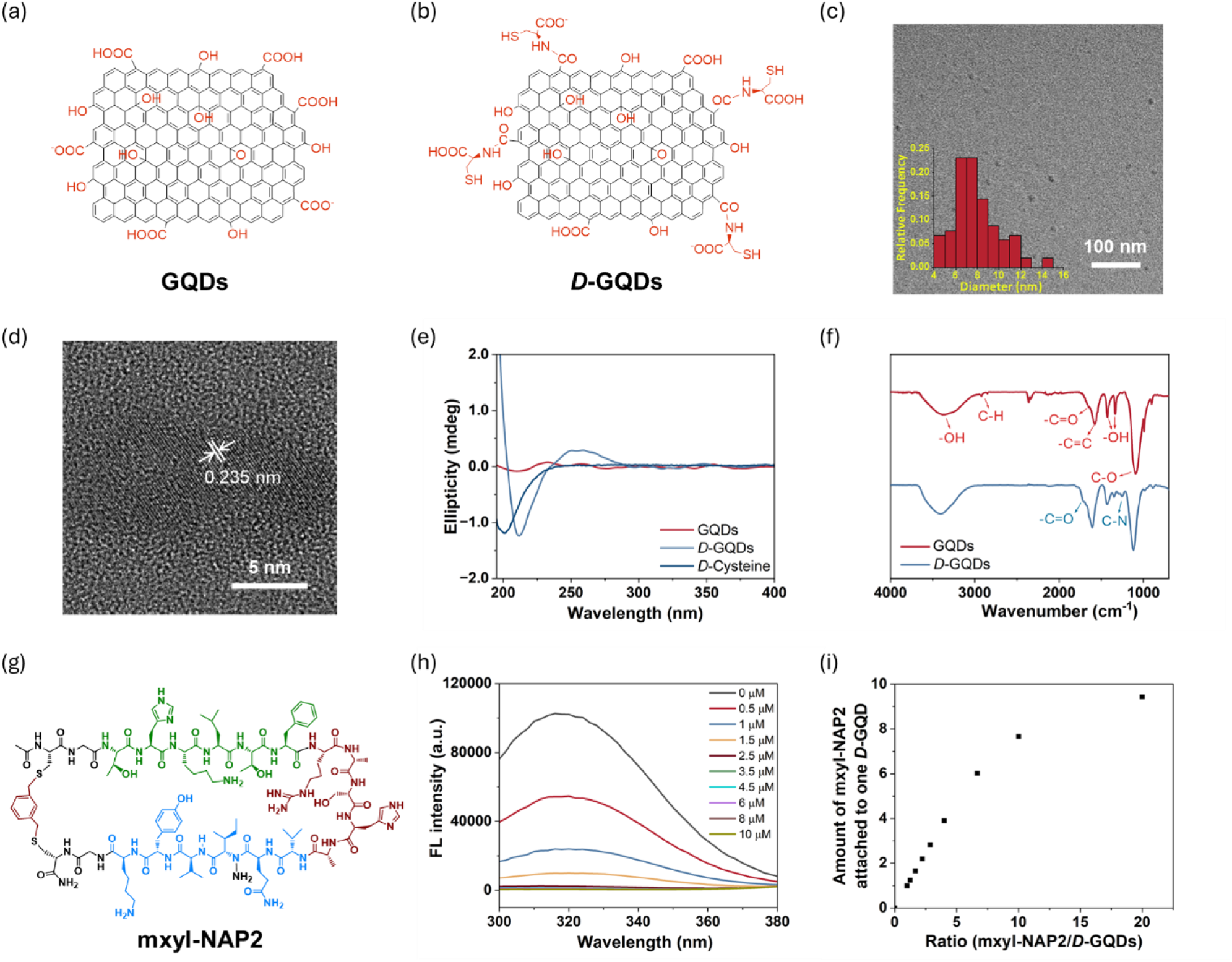
*D*-GQD characterization and its incorporation with tau-targeting mxyl-NAP2. Molecular structure of (a) GQDs and (b) *D*-GQDs. (c-d) Transmission electron microscope (TEM) images and size distribution of *D*-GQDs. (e) Circular dichroism (CD) spectra and (f) Fourier transform infrared (FTIR) spectra of GQDs and *D*-GQDs. (g) Molecular structure of mxyl-NAP2. (h) Fluorescence spectra of mxyl-NAP2 (10 µM) incubated with different concentrations of *D*-GQDs. (i) Amount of mxyl-NAP2 bound to one *D*-GQD by incubation at different ratios of mxyl-NAP2 to *D*-GQDs.

To improve the selectivity of *D*-GQDs towards tau proteins, we incorporated a tau-targeting peptide, mxyl-NAP2, reported in our previous work **(Fig. 2g, Fig. S2)**.^36^ Mxyl-NAP2 was engineered to contain both the aggregation-prone PHF6 sequence and cross-β interaction motifs, inspired by solid-state structure of tau. These design elements allow mxyl-NAP2 to specifically recognize and bind tau proteins with high affinity.

The π-conjugated surface of *D*-GQDs facilitates π-π stacking interactions with the aromatic rings of mxyl-NAP2, reinforcing their binding. Additionally, the interaction is further stabilized by electrostatic attraction between the negatively charged *D*-GQDs and the positively charged mxyl-NAP2 peptides. This dual-binding mechanism ensures robust and selective complex formation, crucial for targeted tau detection. The amount of mxyl-NAP2 bound on each *D*-GQD was determined by the quenching efficiency using the fluorescence resonance energy transfer (FRET) assay **(Fig. 2h)**.^51^ Mxyl-NAP2 showed fluorescence emission centered at 320 nm when excited at 265 nm, due to tyrosine (Tyr) fluorophores in its structure.^52^ Upon titrating increasing concentrations of *D*-GQDs with a fixed 10 µM concentration of mxyl-NAP2, the fluorescence intensity of mxyl-NAP2 decreased, confirming successful binding. At lower peptide-to-*D*-GQD ratios (<7), the amount of bound mxyl-NAP2 increased linearly, indicating efficient surface binding. However, as the ratio exceeded 7, the amount of bound peptide saturated due to the capacity of planar surface on *D*-GQDs **(Fig. 2i)**.

### 2.2. Enhanced selective inhibition of tau aggregation by mxyl-NAP2/*D*-GQD

Upon successful incorporation of mxyl-NAP2 on the *D*-GQD, we investigated whether the tau-targeting peptide can improve the selective inhibition of tau aggregation over abundant proteins in physiological conditions. The tau protein used in these studies includes a P301L mutation (tau_P301L_) which is frequently observed in patients with FTDP-17 **(Fig. S3)**.^53^ Tau_P301L_ contains four microtubule-binding repeat domains (R1–R4), including the PHF6 aggregation-prone motif (VQIVYK, residues 248–253), which can be targeted by mxyl-NAP2.^36, 54^ To assess the selectivity of mxyl-NAP2/*D-*GQD complex to tau, we studied their inhibitory effect on the aggregation of tau_P301L_ with or without the presence of human serum albumin (HSA). The processes were monitored by Thioflavin T (ThT), a benzothiazole dye that exhibits enhanced fluorescence upon binding to amyloid fibrils with β-sheet secondary structure.^55^ HSA is the most abundant circulating protein found in plasma and was reported to interact with GQDs *via* hydrogen bonds and van der Waals forces.^32, 56^ Due to the heparin sulfate in the tau_P301L_ aggregation buffer, HSA was induced to slightly aggregate, resulting in a sigmoidal curve from increased ThT fluorescence **(Fig. S4a)**. *D*-GQDs exhibited inhibitory effect on HSA aggregation due to their interaction with HSA, while mxyl-NAP2 showed negligible effect on its aggregation, indicating its binding specificity **(Fig. S4 b-c)**. In the presence of HSA, *D*-GQDs maintained inhibitory effects on tau_P301L_ aggregation, with efficiencies of 88.9%, 78.6%, and 68.4% at concentrations of 0.4, 0.3, and 0.2 µM respectively, compared to the conditions without HSA **(Fig. 3a)**. Meanwhile, the mxyl-NAP2/*D*-GQD complex (8:1) at higher concentrations retained substantial inhibitory activity against tau_P301L_ aggregation in the presence of HSA. The complex maintained 95.1% and 88.7% of its inhibitory efficiency at 0.4 and 0.3 µM, respectively, compared to the condition without HSA. This indicates that the mxyl-NAP2/*D*-GQDs complex at higher concentrations showed improved selectivity for tau over HSA. Compared to *D*-GQDs, mxyl-NAP2/*D*-GQDs at 0.4 and 0.3 µM also exhibited enhanced inhibitory effect in the presence of HSA by 14.7% and 18.7% due to the inherent inhibition capacity of mxyl-NAP2 to tau aggregation **(Fig. 3b, Fig. S4d)**. At 0.2 µM, mxyl-NAP2/*D*-GQDs did not show improved selectivity for tau nor did it enhance the inhibition efficiency. This result could be attributed to the templating effect of mxyl-NAP2 at a lower concentration, where it led to a slight increase in ThT fluorescence **(Fig. 3a, Fig. S4d)**. We further explored how the amount of mxyl-NAP2 loaded onto the *D*-GQD surface influences the selectivity of the mxyl-NAP2/*D-*GQD complex to tau. We tested the mxyl-NAP2/*D-*GQD complexes at molar ratios of 8:1, 5:1, and 3:1 at a constant *D*-GQD concentration of 0.3 µM, confirmed by the fluorescence quenching assay **(Fig. 2i)**. Among these samples, the 8:1 mxyl-NAP2/*D*-GQD complex demonstrated a significantly enhanced inhibitory efficiency on tau_P301L_ aggregation in the presence of HSA **(Fig. 3c, d)**. This is potentially due to the near-complete surface coverage of the *D*-GQD by mxyl-NAP2, which reduces the interactions with HSA **(Fig. 3e)**. When mxyl-NAP2/*D*-GQD approaches tau proteins, mxyl-NAP2 detaches from *D*-GQDs and form a more stable interaction with tau.^36^ Subsequentially, the exposed *D*-GQDs may further interact with the surrounding tau proteins *via* π-π stacking and electrostatic interactions, thereby enhancing the selectivity of *D*-GQDs for tau.^27, 28^ In contrast, the 5:1 and 3:1 complex did not significantly improve the selectivity of *D*-GQDs for tau **(Fig. 3c, d)**. This is potentially attributed to the π-conjugated planes on the *D*-GQD surface, with an average diameter of around 8 nm, which remains partially exposed at lower mxyl-NAP2 ratios, allowing interactions with HSA **(Fig. 3e)**. The presence of hydrogen bonding between *D*-GQDs and HSA could increase the distance between *D*-GQDs and tau and thus interfere with their interactions, limiting inhibitory efficiency on tau aggregation.^32^ Overall, the results suggested that at concentrations of 0.4 and 0.3 µM, the 8:1 mxyl-NAP2/*D*-GQD complex significantly improved the selectivity of *D*-GQDs to tau over HSA. This improved selectivity is attributed to the specific affinity of mxyl-NAP2 for fibrillar tau.

**Fig. 3.**
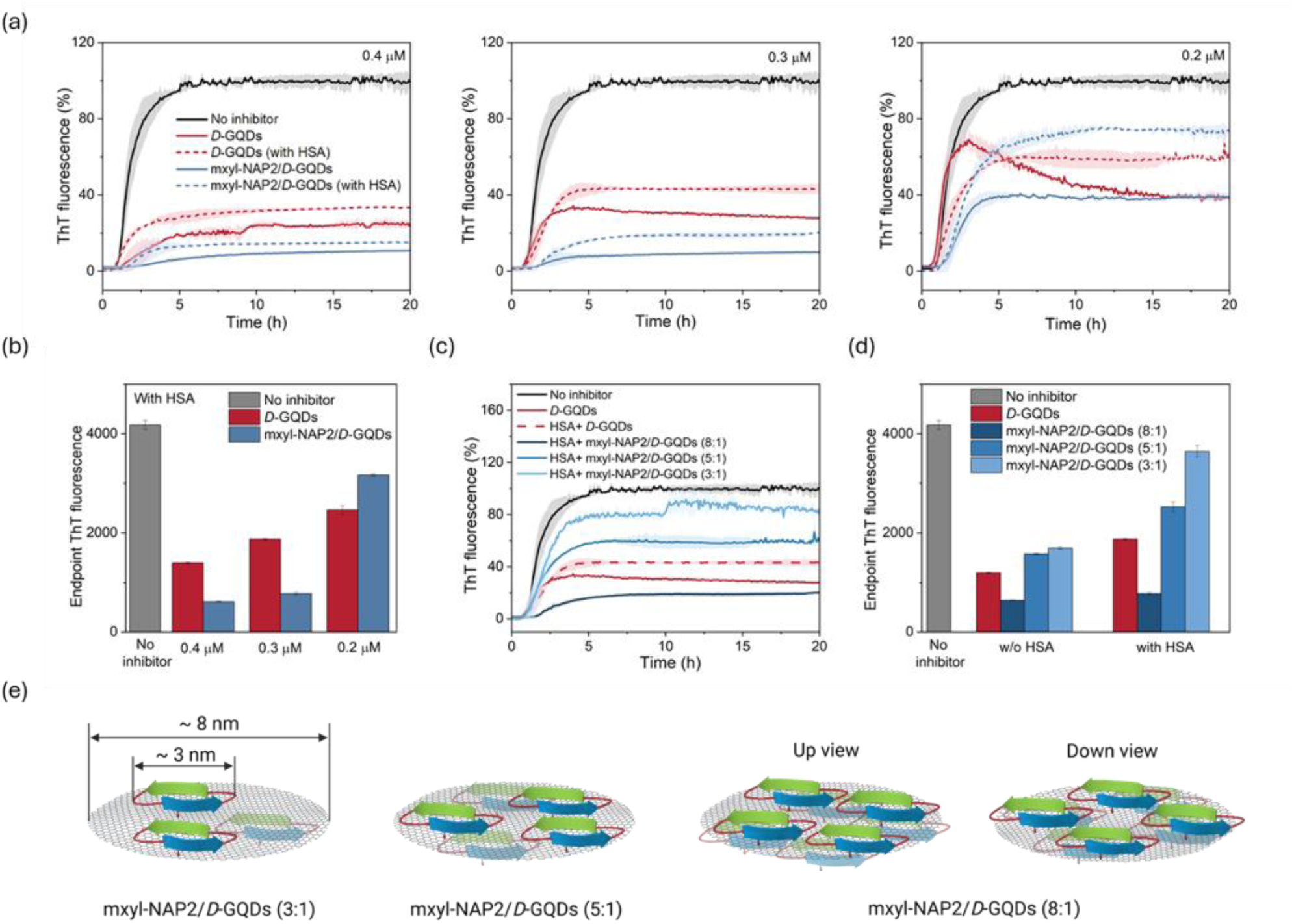
(a-b) Thioflavin T (ThT) fluorescence of tau_P301L_ incubated with 0.4, 0.3, and 0.2 µM of *D*-GQDs or mxyl-NAP2/*D*-GQD complex (mxyl-NAP2: *D*-GQDs= 8:1) in the presence and absence of human serum albumin (HSA). (c-d) ThT fluorescence of tau_P301L_ incubated with 0.3 µM of *D*-GQDs or mxyl-NAP2/*D*-GQD complex (mxyl-NAP2: *D*-GQDs= 8:1, 5:1, and 3:1) in the presence and absence of HSA. Fluorescent signals of all samples were normalized to the control (without inhibitors). (e) Schematic illustration of mxyl-NAP2/*D*-GQD complex at different molar ratios: Mxyl-NAP2 interacted with *D*-GQD π-conjugated plane *via* π-π stacking.

### 2.3. Efficient encapsulation of mxyl-NAP2/*D*-GQDs in sEVs

To further improve the off-target effect of *D*-GQDs, we encapsulated the mxyl-NAP2/*D*-GQD complex into sEVs, utilizing the highly efficient *D-*GQD loading technique developed in our prior work.^46^ sEVs were isolated from cell culture media of 3T3 cells, a mouse fibroblast cell line, using size-based ultrafiltration techniques.^23^ They displayed a spherical shape with structural integrity, confirmed by TEM, and had an average diameter of 127.2 ± 1.8 nm, measured by nanoparticle tracking analysis (NTA) **(Fig. 4a, b)**. Moreover, western blot analysis of sEV lysates confirmed the expression of sEV biomarkers. The results showed clear immunoblotted bands for CD9, CD63, and CD81, which are commonly enriched in the sEV membrane, suggesting that the isolated nanoparticles were sEVs as expected **(Fig. 4c)**.^57^ To encapsulate mxyl-NAP2/*D-*GQD complex into sEVs, 8 µM of the complex was incubated with 3T3 sEVs (1×10^9^ particles/mL) in phosphate-buffered saline (PBS) buffer (pH 7.4) at room temperature under a static condition. Following the incubation, we verified the size and concentration of loaded sEVs by NTA **(Fig. S5)**, and encapsulation of the mxyl-NAP2/*D*-GQDs complex into sEVs was confirmed by fluorescent confocal microscopy. In brief, the sEVs were labeled with a red fluorescence lipid membrane dye (DiI), and mxyl-NAP2/*D*-GQD complex was tracked by the intrinsic blue fluorescence emission of *D*-GQDs excited at 365 nm **(Fig. 4d)**. The colocalization of DiI-labeled sEVs (red) and *D*-GQDs (blue) indicated the permeation and accumulation of the mxyl-NAP2/*D*-GQD complex inside sEVs. Furthermore, we determined that the permeation efficiency of *D*-GQDs in the mxyl-NAP2/*D*-GQD loaded sEVs was 65.7 ± 4.2% by the quantification of the *D*-GQD blue signals within sEVs (See Method).^46^ To determine the encapsulation efficiency of mxyl-NAP2 loaded by *D*-GQDs in sEVs, mxyl-NAP2 was separated from *D*- GQDs and other components in the lysis of the loaded sEV by using a 2-kDa centrifuge tube. The concentration of encapsulated mxyl-NAP2 was confirmed based on a colorimetric assay. The UV-vis absorbance spectrum of mxyl-NAP2 showed a distinct absorption peak at 292 nm, which corresponds to the presence of tyrosine in its structure.^52^ The absorbance demonstrated a linear relationship with concentration **(Fig. 4e, f)**. Using this calibration curve, the concentrations of encapsulated mxyl-NAP2 were determined to be 24.67, 18.50, and 10.08 µM for sEVs loaded with mxyl-NAP2/*D-*GQD ratios of 8:1, 5:1, and 3:1 **(Fig. 4g)**. Thus, the encapsulation efficiencies of mxyl-NAP2 into sEVs were 30.8%, 38.5%, and 42%, respectively **(Table S1)**.

**Fig. 4.**
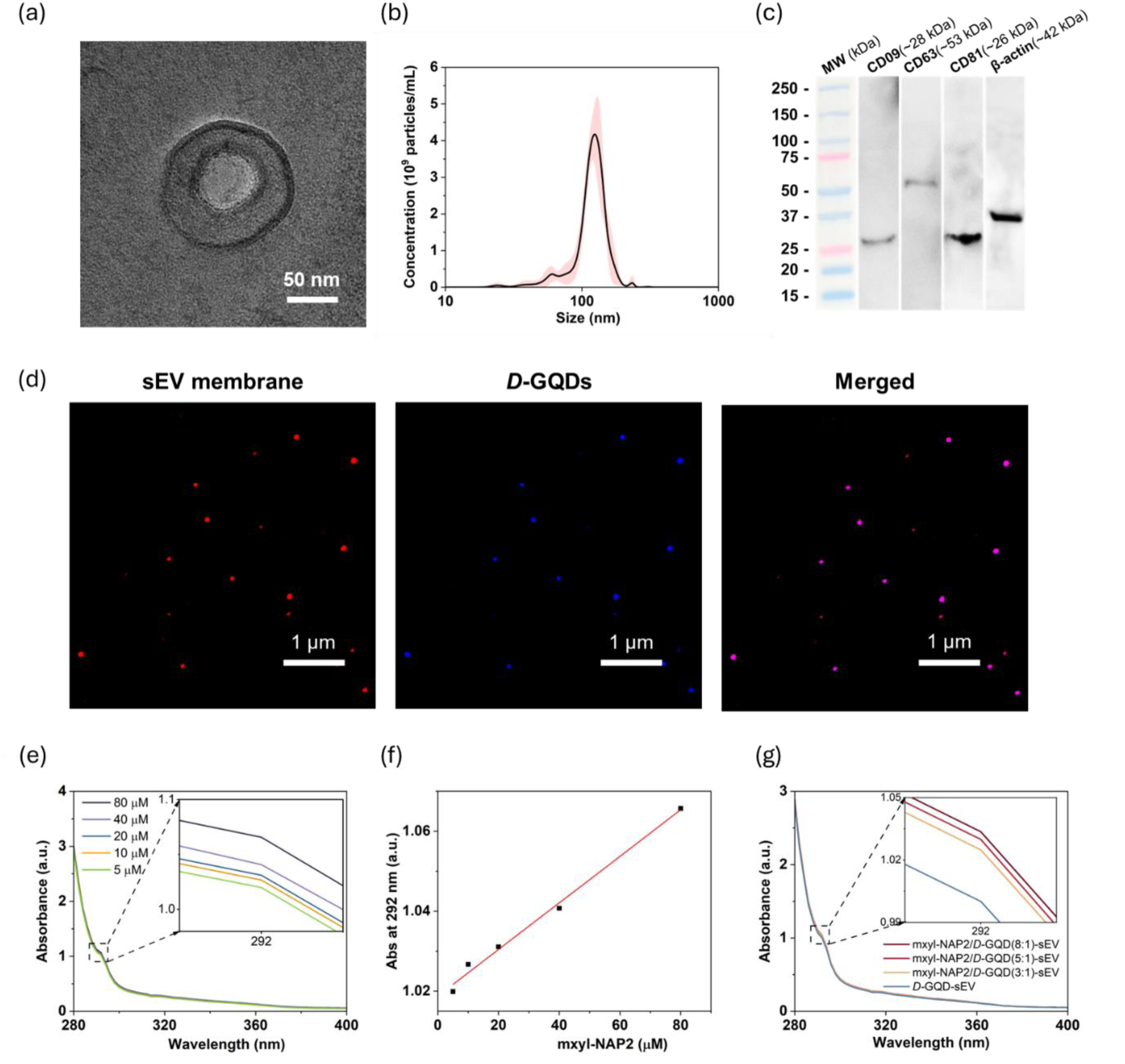
(a) TEM image and (b) nanoparticle tracking analysis (NTA) of small extracellular vesicles (sEVs). (c) Western blot analysis of sEV biomarkers. (d) Confocal Laser Scanning Microscopy (CLSM) images of mxyl-NAP2/*D*-GQDs (blue) loaded DiI-labeled sEVs (red). (e-f) The UV-vis absorbance spectra of mxyl-NAP2 at different concentrations in the presence of a constant concentration of Tween-20. The absorbance at 292 nm exhibited a linear relationship with its concentration. (g) The UV-vis absorbance spectra of encapsulated mxyl-NAP2 into sEVs, which was separated from *D*-GQDs and sEV membrane after lysis with surfactant Tween-20, with different ratios of mxyl-NAP2 to *D*-GQDs (8:1, 5:1, and 3:1).

### 2.4. Effective inhibition of tau propagation by mxyl-NAP2/*D*-GQD loaded sEVs

A key feature of tau NFTs is their ability to propagate between neurons, spreading tau pathology from diseased cells to adjacent healthy cells.^58, 59^ In previous studies, we demonstrated that *D*-GQDs and mxyl-NAP2 could inhibit the seeding activity of tau fibrils respectively using the cellular tau biosensor assay. To quantify the number of intracellular aggregates and evaluate whether inhibitors can prevent the cellular transmission of tau fibrils, we employed HEK293 biosensor cells that stably expressed a tau-yellow fluorescent protein fusion [tau RD(LM)-YFP] **(Fig. 5a)**.^9, 60^ The aggregation of endogenous tau induced by extracellular seeds led to focal puncta with green fluorescence, which can be quantified by brightness threshold-based analysis of green puncta. The results showed a significant reduction in the seeding activity of tau fibrils, decreasing from 71% to 34% as the concentration of *D*-GQDs increased from 0.05 µM to 0.3 µM, compared to the control group without inhibitors **(Fig. 5b-c, Fig. S7)**. Notably, *D-*GQD-sEV exhibited significantly higher inhibitory efficiency on tau fibrils seeding activity than *D*-GQDs at the same concentration range. This improvement is potentially attributed to the ability of sEVs to shield *D*-GQDs from interacting with other biomolecules in the cell media such as bovine serum albumin, thereby enhancing the bioavailability of *D*-GQDs in cells. Moreover, as the ratio of mxyl-NAP2 to *D*-GQDs increased, the seeding activity of tau fibrils was further decreased. Remarkably, when the mxyl-NAP2/*D-* GQDs complex encapsulated in sEVs were used at a concentration of 0.3 µM, the inhibitory efficiency of mxyl-NAP2/*D*-GQD (8:1)-loaded sEVs was enhanced by 9.4% compared to *D*-GQD-sEVs alone. To note, mxyl-NAP2 alone showed low efficacy when added directly to cells due to limited cellular uptake **(Fig. S8)**. These findings are consistent with previous ThT assay results, suggesting that higher levels of mxyl-NAP2 bound to the surface of *D*-GQDs improve their selectivity for tau fibrils.

**Fig. 5.**
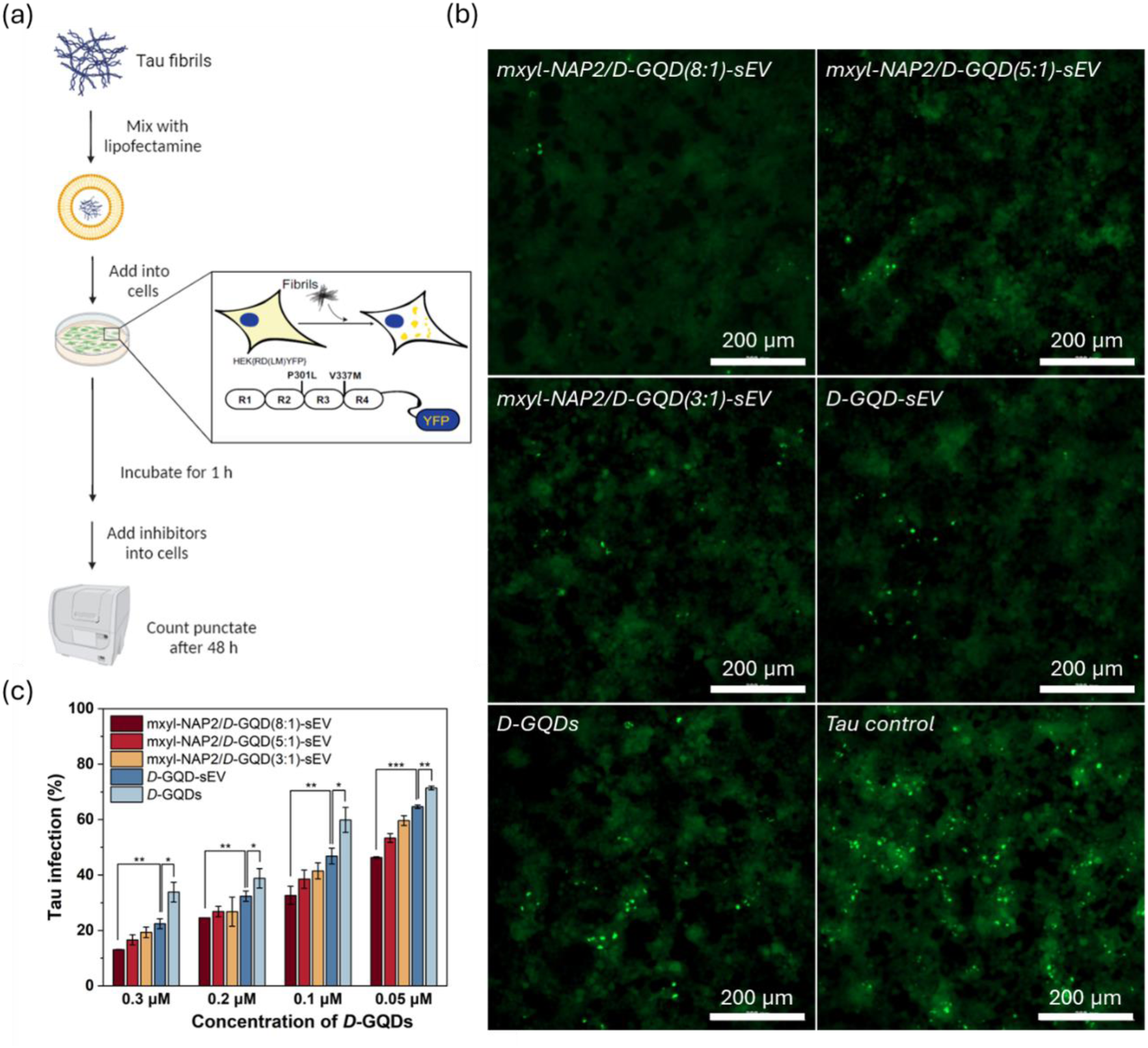
(a) Scheme of the workflow of cellular tau biosensor propagation assay. (b) Mxyl-NAP2/*D*-GQD-sEVs, *D-* GQD-sEVs, and *D*-GQDs, at a concentration of 0.3 µM for *D*-GQDs, sEV (1×10^9^ particles/mL), prevented the cellular transmission of mature tau fibrils (0.19 µM). Fluorescent images of cellular tau biosensors were taken under the FITC channel (ex/em: 469/525 nm). The green puncta with high fluorescence represented the tau aggregation in cells induced by exogenous tau_P301L_ fibers. Scale bars: 200 μm. (c) Tau infection (%) in the column graphs showed the number of intracellular fluorescent puncta, normalized to cells treated with tau fibrils only.

### 2.5. Enhanced inhibition of tau propagation between neurons *via* RVG-modified sEVs

Since tau proteins are predominantly found in neurons, it is crucial to enhance the neuronal targeting ability of the developed inhibitors to neurons. To achieve this, we modified the surface of sEVs with a 29-amino-acid peptide, derived from RVG. This peptide specifically interacts with the nicotinic acetylcholine receptor (nAchR) to enable viral entry into neuronal cells, which has been utilized to target neurons in drug delivery systems.^61–63^ To examine whether RVG modified sEVs can further improve inhibitory efficiency on the seeding activity of tau fibrils between neurons, we used thioflavin S (ThS) staining to visualize intracellular tau aggregates. ThS was chosen due to its strong green fluorescence when bound to amyloid-like aggregates with β-sheet secondary structures, allowing clear observation of tau accumulation within the cells.^64^ In parallel, we employed immunofluorescence with monoclonal antibody Tau46 to confirm the tau expression in SH-SY5Y cells.^65^ In this setup, tau fibrils were introduced into SH-SY5Y cells one hour prior to the addition of the inhibitors. After 24 hours of incubation, intracellular tau aggregates were visualized using confocal microscopy. The extent of tau aggregation was quantified by measuring the green fluorescence pixel intensity within the cells. The results demonstrated that both RVG-*D*-GQD-sEV and RVG-mxyl-NAP2/*D*-GQD-loaded sEV significantly reduced tau fibril seeding activity between neurons at concentrations ranging from 0.05 µM to 0.2 µM, compared to *D*-GQD-sEV and mxyl-NAP2/*D*-GQD-loaded sEV **(Fig. 6, Fig. S9)**. To better understand this phenomenon, we examined the cellular uptake of sEV and RVG-sEV in SH-SY5Y cells. The results showed that the neuronal uptake of RVG-sEV was 2.3 times higher than that of unmodified sEV after 4-h treatment. This was quantified by measuring the pixel intensity of sEV stained with DiI (a red fluorescent dye for lipids) within the cells stained with green cytoplasmatic membrane dye **(Fig. S10)**. The increased uptake efficiency can be attributed to RVG-sEV’s ability to selectively bind to nAchR expressed on neurons, thereby increasing the binding affinity and facilitating more effective neuronal entry.^62, 66^ The enhanced uptake of RVG-sEV led to more efficient intracellular delivery of tau inhibitors, resulting in a significant improvement in inhibiting tau transmission between neurons.^66–68^

**Fig. 6.**
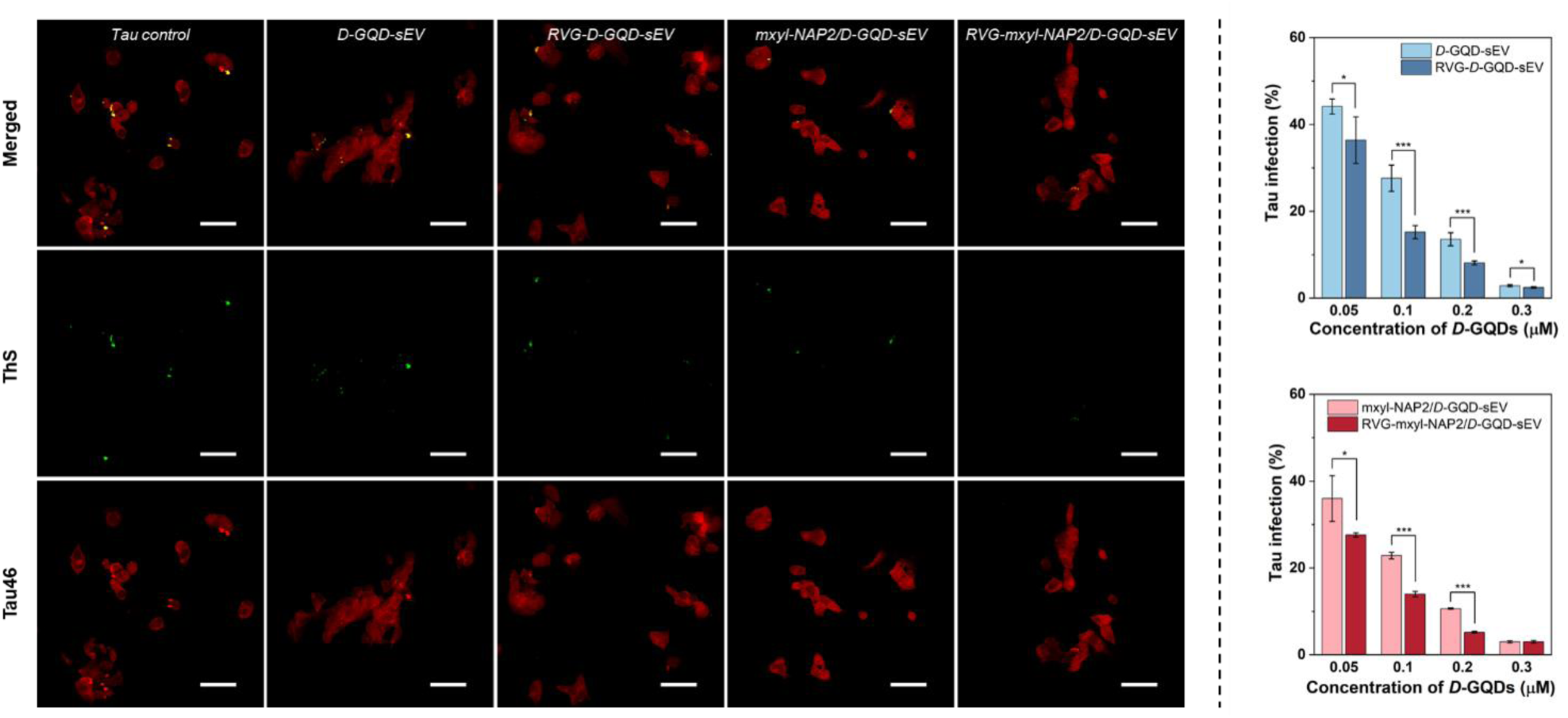
(a) CLSM images of tau fibril-treated SH-SY5Y cells incubated with different inhibitors at a concentration of 0.05 µM for *D*-GQDs. SH-SY5Y cells were sequentially double-stained with antibody Tau46 (red) for intracellular tau and Thioflavin S (ThS, green) for tau fibrils. Scale bar: 50 µm. Bar graphs show the pixel number of tau fibrils in the green channel per cell relative to control without inhibitors.

## 3. Conclusion

This study demonstrates the development of a RVG-mxyl-NAP2/*D*-GQD-loaded sEV drug delivery system as a promising, targeted therapeutic approach for tauopathies. By integrating mxyl-NAP2—a tau-specific binder—onto *D*-GQD and encapsulating mxyl-NAP2/*D*-GQD in sEVs, we achieved greater selectivity of *D*-GQD for tau, reducing off-target interactions with non-target proteins in physiological conditions. Additionally, the neuron-targeting capability provided by RVG-modified sEVs facilitated the efficient delivery of mxyl-NAP2/*D*-GQD to neuronal cells, significantly enhancing the inhibition of tau aggregation and propagation. This targeted system addresses the limitation of GQD inhibitors for tau proteins, offering a robust and selective therapeutic platform for treating tauopathies with reduced off-target effects and improved therapeutic efficacy against neurodegenerative diseases. Future work will further investigate the clinical translatability of this approach, particularly regarding long-term safety and efficacy in relevant *in vivo* models.

## 4. Experimental Section

### Materials

Sulfuric acid (95–98%) and nitric acid (69–70%) were purchased from VWR (PA, USA). Carbon nanofibers, sodium hydroxide, N-hydroxysulfosuccinimide sodium salt (Sulfo-NHS, >98%), sodium acetate (>99%), ThT, heparin sodium salt, and DL-dithiothreitol (DTT, 97%), HSA, and ThS were purchased from Sigma-Aldrich (MO, USA). The dialysis membrane tubing (MWCO: 1kD) was purchased from Spectrum Chemical Manufacturing Company (NJ, USA). 1-Ethyl-3-(3-dimethylaminopropyl) carbodiimide (EDC), DMEM medium, Opti-MEM medium, DMEM/F12 medium, fetal bovine serum (FBS), Lipofectamine 2000 transfection reagent, bovine serum albumin (BSA, 10%), and antibiotic-antimycotic were purchased from Thermo Fisher Scientific (MA, USA). *D*-Cysteine was purchased from AmBeed (IL, USA). Tween-20 (10%) was purchased from G-Biosciences (MO, USA). Paraformaldehyde (4%) was ordered from Electron Microscopy Sciences (PA, USA). Triton X-100 (25%) was purchased from GeneTex (CA, USA). Phosphate-buffered saline (PBS) was purchased from Corning (NY, USA).

### Synthesis and characterization of GQDs and *D*-GQDs

GQDs were synthesized using a modified Hummers’ method.^48^ 0.45 g of carbon fiber was added into 90 mL of concentrated H_2_SO_4_ (98%) and stirred for 1.5 h. Then, 30 mL of concentrated HNO_3_ (68%) was added into the mixture solution and sonicated for 1 h. The mixture reacted at 120 °C for 20 h. Next, the solution was neutralized using a sodium hydroxide solution, and further purified through dialysis for 3 days with a dialysis bag (retained molecular weight: 1000 Da). The synthesis of *D*-GQDs was carried out using EDC/NHS coupling reaction.^69^ 1 mL of EDC (100 mM) was added into 25 mL of GQDs (12.5 µM) and stirred for 30 min. 1 mL of NHS (500 mM) was added into the mixture solution and stirred for another 30 min. Then, 1 mL of *D*-cysteine (100 mM) was added into the reaction and the mixture was reacted for 16 h. The product was purified using a 1 kDa MWCO dialysis bag.

The size and shape of *D*-GQDs were characterized by TEM (Thermo Scientific Talos F200i; MA, USA) under an accelerating voltage of 200 kV. A 3 µL droplet of the *D*-GQD solution (5 µM) was placed on the carbon-coated copper TEM grid (Purchased from Electron Microscopy Sciences; PA, USA), and air-dried. The size distribution of the *D*-GQDs was analyzed using ImageJ software. The chemical composition was performed by ATR-FTIR using Bruker Tensor 27 FTIR Spectrometer (Bruker Optics Inc., MA, USA) with a diamond lens Attenuated Total Reflectance (ATR) module. 3 µL of 100 µM GQDs or *D*-GQDs was dried in the air and each spectrum was measured as the accumulation of 64 scans at a spectral resolution of 2 cm^-1^ within the range 4000–700 cm^-1^. The fluorescence emission property of GQDs and *D*-GQDs was measured using a plate reader (Tecan Infinite 200Pro; Männedorf, Switzerland). The fluorescence was measured at an excitation wavelength of 365 nm and an emission wavelength from 400 nm to 700 nm. The zeta potential of GQDs and *D*-GQDs (6.25 µM) was detected using Malvern Zetasizer Nano ZS (Malvern Panalytical; Worcestershire, United Kingdom) with results averaged over three runs. The chiroptical activity and absorption of GQDs and *D*-GQDs were tested by CD spectroscopy (Jasco J-1700 Spectrometer; MD, USA). Samples were diluted to 2.5 µM and scanned from 200 nm to 400 nm with 0.1 nm intervals, 5 nm bandwidth, and a scan speed of 50 nm min^-1^.

### Synthesis of mxyl-NAP2

^36^ Automated solid-phase peptide synthesis was carried out on NovaPEG Rink amide resin (35-100 mesh, 0.45 mmol/g). Linear precursors for macrocyclization were synthesized at 0.25 mmol scale. The following amino acid derivatives suitable for Fmoc SPPS were used: Fmoc-Asp(tBu)-OH, Fmoc-Glu(tBu)-OH, Fmoc-Arg(Pbf)-OH, Fmoc-Gly-OH, Fmoc-Cys(Trt)-OH, Fmoc-Thr(tBu)-OH, Fmoc-His(Boc)-OH, Fmoc-Leu-OH, Fmoc-Phe-OH, Fmoc-Ser(tBu)-OH, Fmoc-D-Ala-OH, Fmoc-Gln(Trt)-OH, Fmoc-Asn(Trt)-OH, Fmoc-Tyr(tBu)-OH, Fmoc-Lys(Boc)-OH, Fmoc-Val-OH, Fmoc-Ile-OH, and Fmoc-Gln(Trt)-(Boc)aIle-OH that was synthesized previously. Fmoc deprotection steps were performed by treating the resin with a solution of 20% piperidine/DMF once at room temperature (5 min), and then at 75°C (2 min). Following Fmoc deprotection the resin was washed 4× with DMF. Coupling of Fmoc-protected amino acids was performed using 5 equiv HCTU (0.25 M in DMF), 10 equiv NMM (1 M in DMF), and 5 equiv of Fmoc-protected amino acid or dipeptide (0.2 M in DMF) at 50°C (10 min × 2). Deprotection and coupling steps were repeated until peptide synthesis was complete and then a final Fmoc deprotection was run to remove Fmoc from the N-terminus. The resin was transferred to a suitable vessel, washed with DCM (5 mL × 4) and dried under vacuum. Peptides were N-terminally acetylated using 5% acetic anhydride and 10% pyridine in DCM (15 min), washed with DCM (5 mL × 4), and dried under vacuum before being cleaved from the solid support and globally deprotected by incubating the dried resin in 4 mL of TFA:TIPS:H_2_O:DODT (92.5:2.5:2.5:2.5) for 2.5 h. The resin was filtered, and the filtrate was collected in a 50 mL centrifuge tube. The resin was washed with DCM (10 mL) and filtered, and crude peptides were precipitated from the combined filtrate by the addition of cold Et_2_O (40 mL). The mixture was centrifuged and the supernatant was decanted. The pellet was washed with Et_2_O (25 mL × 2) and dried thoroughly under vacuum. For dithiol bis-alkylation cyclization, 1.5 equiv of 1,3-bis(bromomethyl)benzene was added to a 1 mM solution of linear precursor peptide in 1:1 MeCN:H_2_O buffered with NH_4_HCO_3_ (20 mM) and the pH was adjusted to 8.0 using 2 M aq NaOH. Reaction progress was monitored by analytical HPLC. The reaction was stirred for 2 h before evaporating the MeCN under a stream of N_2_, freezing, and lyophilization. Mxyl-NAP2 was purified by preparative RP-HPLC (C12, 250 mm × 21.2 mm, 4 μm, 90 Å) using linear gradients of MeCN in H_2_O (mobile phases modified with 0.1% formic acid) over 30 minutes. Analytical RP-HPLC spectra were acquired (C12, 150 mm × 4.6 mm, 4 μm, 90 Å) using linear gradients of MeCN in H_2_O (mobile phases modified with 0.1% formic acid) over 20 minutes. HRMS was acquired using a Bruker Impact II ESI-QTOF.

### Tau_P301L_ expression and purification

Human tau_P301L_ (0N4R) with an N-terminal His_6_ tag was purified following the previous protocol with a slight modification.^70^ Briefly, transformed BL21 (DE3) cells were cultured in LB + Kanamycin media at 37°C until OD_600_ reached between 0.6 - 0.8 and were then induced with 0.5 mM IPTG overnight at 16 °C. Cells were then harvested, resuspended, and lysed by probe sonication in the lysis buffer containing 20 mM Tris, 500 mM NaCl, 10 mM imidazole, and 5 mM serine protease inhibitor PMSF, adjusted to pH 8.0. The lysate was then boiled in a water bath for 20 minutes and the debris was pelleted by centrifugation at 20,000g for about 40 min at 4 °C. The resulting supernatant was injected into a 5 mL IMAC Ni-Charged affinity column and eluted over a gradient of 10–200 mM imidazole. Eluted tau-containing fractions were further purified using GE HiPrep 16/60 Sephacryl S-200 high-resolution size exclusion chromatography into a storage buffer containing 20 mM Tris, 150 mM NaCl, and 1 mM DTT, adjusted to pH 7.6. The purity of the protein was confirmed by sodium dodecyl sulfate-polyacrylamide gel electrophoresis (SDS-PAGE) analysis, and the concentration was determined using bicinchoninic acid (BCA) assay.

### Fluorescence Resonance Energy Transfer (FRET) Assay

10 µM of mxyl-NAP2 in PBS was incubated with different concentrations of *D*-GQDs at room temperature in a 96-well plate. After 10-min incubation, the fluorescence of the mixture solution was measured at an excitation wavelength of 265 nm and an emission wavelength from 300 nm to 380 nm using a Tecan Infinite 200 Pro plate reader. Background fluorescence from the *D*-GQD solution was subtracted from all measurements. The amount of mxyl-NAP2 attached to one D-GQD was calculated using the following formula:

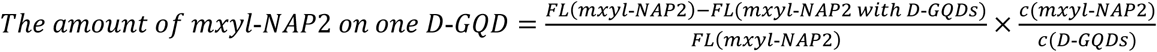

### ThT Aggregation Assay

Recombinant tau_P301L_ (5 µM final concentration), human serum albumin (HSA, 5 µM final concentration), and *D*-GQDs or mxyl-NAP2/*D*-GQD complex with molar ratios of 8:1, 5:1, and 3:1 (final concentration of *D*-GQDs: 0.4, 0.3, and 0.2 µM) were mixed in an aggregation buffer (100 mM sodium acetate, 10 µM ThT, 5 µM heparin, and 2 mM DTT, pH: 7.4). 200 µL of the same solution was added into two wells of a 96-well plate for each sample. The plate was then sealed with a clear sealing film and allowed to incubate at 37 °C in a Tecan Infinite 200Pro plate reader. ThT fluorescence measurements were conducted automatically after 30 s shaking at an excitation wavelength of 450 nm and an emission wavelength of 485 nm at an interval of every 5 min for 20 h. Every experiment included control wells that lacked tau_P301L_, HSA, or inhibitors.

### Isolation and Characterization of sEVs

When 3T3 cells reached 70-80% confluency in the flask, cell-culture medium (CM; MEM with 10% FBS and 1% Antibiotic-Antimyotic) was replaced into serum-free CM after washing five times with PBS. The serum-free CM was collected after 24 h. Then, the serum-free CM was filtered with Pore Size 0.22 µm Vacuum Filtration Systems (VWR, PA, USA) to remove undesired large debris. Then, the filtered CM was washed with PBS buffer and concentrated using a 100 kDa centrifuge tube (Spin-X UF Concentrator; Corning, NY, USA). The morphology and nanostructure of sEVs were observed by TEM (JEOL 2011, JEOL Ltd., Tokyo, Japan) under an accelerating voltage of 120 kV with negative staining by UranyLess. The size distribution and concentration of sEVs were determined with a NanoSight NS300 (NanoSight Ltd., UK) with a 1000-fold dilution of sEVs and analyzed with NTA 3.3 analytical software suite. sEV marker expressions (CD9, CD63, and CD81) were validated through western blot. Briefly, 15 µg of proteins of sEV lysates were separated by SDS-PAGE. A nitrocellulose membrane (Bio-Rad Laboratories, CA, USA) was used to transfer the separated proteins to SDS gel. Next, the primary antibodies (anti-beta-actin, anti-CD9, anti-CD63, and anti-CD81; Santa Cruz Biotechnology, Inc., TX, USA) were incubated overnight with the transferred membrane followed by secondary antibodies (anti-HRP-linked secondary antibody; Santa Cruz Biotechnology, Inc., TX, USA) were then blotted. The immunoreactive species were detected by an enhanced chemiluminescence (ECL) substrate using a C400 Bioanalytical Imager (Azure Biosystems, CA, USA).

### Permeation of *D*-GQDs into sEVs

8 µM of *D*-GQDs were incubated with sEVs (final concentration: 1×10^9^ particles/mL) at room temperature for 20 min to ensure full permeation. sEV membranes were stained with DiI dye (Biotium, CA, USA) for imaging with confocal laser scanning microscopy (CLSM; A1R-MP Laser Scanning Confocal Microscopy, Nikon, Tokyo, Japan; ex/em: 561/570-620 nm). Then, sEVs were washed with PBS (8 °C) three times with a 100 kDa centrifuge tube to remove unincorporated free *D*-GQDs. The permeation efficiency of *D*-GQDs into sEVs was indirectly determined by statistically analyzing the count of blue-fluorescent lit-up sEVs (caused by permeation) under confocal microscopy over the total concentration of sEV, which was developed in our previous study. Briefly, 5 μL of loaded sEV sample was evenly spread on an 18 mm × 18 mm cover glass (Corning, NY) and imaged by CLSM with DAPI channel (ex/em: 405/425-525 nm) under a 100× objective. Four random regions were selected, and Z-stack images were captured. Each Z-stack contained 30 images from presenting to disappearing fluorescent dots with a step size of 0.125 μm. Then, captured images were analyzed using ImageJ. The total fluorescent sEV particles (TFEPs) were quantified by settings with manually adjusted thresholds and matching the size of sEVs. Colocalized fluorescent sEV particles (CFEPs) of z-stack images, between successive image sets, were counted *via* ImageJ software with JACOPx Plugin. The total sEV concentration (TEC, particles/mL) was confirmed by NTA. The permeation efficiency of *D*-GQDs into sEVs was calculated using the following formula:

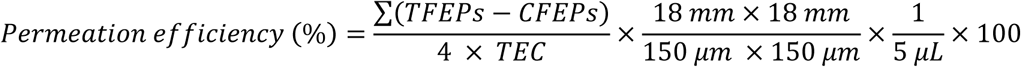

### Determination of Encapsulation Efficiency of mxyl-NAP2 in sEVs

mxyl-NAP2 was incubated with *D*-GQDs at molar ratios of 10:1, 6:1, and 3:1 for 20 min to attach to *D*-GQD surface. To load the complex into sEV, purified sEV (final concentration: 1×10^9^ particles/mL) was gently mixed with mxyl-NAP2/*D*-GQD complex (final concentration of *D*-GQDs: 8 µM) in 200 µL of PBS, and the mixture was incubated at room temperature for 20 min to ensure the full permeation. sEVs were then washed with PBS (8 °C) three times under the support of a 100 kDa centrifuge tube to remove the unincorporated free mxyl-NAP2/*D*-GQD complex. Then, 10 µL of biocompatible surfactants, Tween-20, was added into mxyl-NAP2/*D*-GQD-sEV to lyse the sEV membrane and destroy the interaction between mxyl-NAP2 and *D*-GQDs. To separate mxyl-NAP2 from *D*-GQDs and sEVs, mxyl-NAP2 was washed out in the eluent with a 2kDa centrifuge tube (Sartorius, Germany) at 12.3k rpm for 30 min. The concentration of mxyl-NAP2 in the eluent was quantified by detecting their absorbance at 292 nm using a Tecan infinite 200Pro plate reader. The encapsulation efficiency of mxyl-NAP2 was calculated using the following formula:

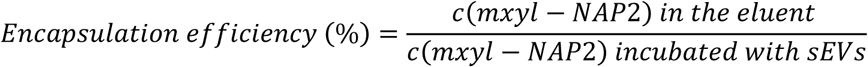

### Cellular Seeding Assay

HEK293 cells stably expressing tau-RD (LM)-YFP were cultured in DMEM complete medium containing 10% FBS, 1% penicillin/streptomycin, and 1% Glutamax under 5% CO_2_ at 37 °C. 90 µL of cells were seeded at a density of 15,000 cells per well in a 96-well tissue culture plate and incubated overnight. To prepare mature tau fibrils for cell seeding, the tau_P301L_ was diluted to a final concentration of 10 µM in an aggregation buffer containing 100 mM sodium acetate, 10 µM heparin, and 2 mM DTT, pH: 7.4. The protein was incubated for 4 days at 37 °C. Following incubation, 8 µL of tau aggregation solution was mixed with 32 µL of low-serum Opti-MEM medium and 2 µL of Lipofectamine 2000. The mixture solution was incubated at room temperature for 20 min. Then, 10 µL of mixture solution was added to HEK-293 cells.

After 1 h adding tau fibrils, 10 μL of inhibitors were added to HEK-293 cells with *D*-GQDs final concentrations around 0.3, 0.2, 0.1, and 0.05 µM in cells and incubated for 48 h at 37 °C. Inhibitors included *D*-GQDs, *D*-GQD-sEV, and mxyl-NAP2/*D*-GQD-loaded sEV. Briefly, mxyl-NAP2 was incubated with *D*-GQDs at molar ratios of 10:1, 6:1, and 3:1 for 20 min to attach to *D*-GQD surface. To load the complex into sEV, purified sEV (final concentration: 1×10^9^ particles/mL) was gently mixed with mxyl-NAP2/*D*-GQD complex (final concentration of *D*-GQDs: 8 µM) in 200 µL of PBS, and the mixture was incubated at room temperature for 20 min to ensure the full permeation. sEVs were then washed with PBS (8 °C) three times under the support of a 100 kDa centrifuge tube to remove the unincorporated free mxyl-NAP2/*D*-GQD complex. The concentration of *D*-GQDs loaded into sEVs was quantified by detecting their intrinsic fluorescence at 520 nm with excitation at 365 nm using a Tecan infinite 200Pro plate reader **(Fig. S6)**.

Every experiment included control wells that lacked tau_P301L_ or inhibitors. After incubation, cells were imaged by a BioTek Cytation 5 cell imager and a microplate reader. 10 × 10 pictures per well were taken at 20× magnification under a fluorescein isothiocyanate (FITC) channel (ex: 469 nm/em: 525 nm), and the punctate counting was carried out using built-in software.

### sEV Modification with RVG Peptide

The sEVs were modified with a 29-amino-acid peptide (YTIWMPENPRPGTPCDIFTNSRGKRASNG; GenScript USA, Inc., NJ, USA), derived from RVG using the previous method.^71^ DOPE-PEG_3400_-NHS (dioleoylphosphatidylethanolamine poly (ethylene glycol)_3400_ N-hydroxysuccinimide; Nanocs Inc., NY, USA) and RVG peptide were combined and allowed to react for 1 h to obtain the DOPE-PEG_3400_-RVG. The DOPE-PEG_3400_-RVG was then incubated with the sEVs with a lipid: sEV ratio of 600:1 for 10 min at 37 °C considering the length of the peptide and radius of sEV. sEVs were then washed with PBS three times (8 °C) to remove the excess DOPE-PEG_3400_-RVG through the use of a 100 kDa centrifuge tube.

### Cellular Uptake of RVG-sEVs

SH-SY5Y human neuroblastoma cells were cultured in the DMEM/F12 complete medium containing 10% FBS and 1% penicillin/streptomycin under 5% CO_2_ at 37 °C. SH-SY5Y cells were seeded at a density of 500,000 cells in the glass bottom cell culture dish (Wuxi NEST Biotechnology Co., Ltd., China) and incubated overnight. sEV were stained by DiI (Biotium, CA, USA), a red fluorescent lipid membrane dye, before adding into cells. sEV and RVG-sEV were incubated with cells (800 sEV/cell) for 4 h. After incubation, the cells were washed with PBS three times to remove residual sEVs, and fixed with 2% paraformaldehyde for 20 min. Then, the cellular membrane was stained by CellBrite Green Cytoplasmic Membrane Dye (Biotium, CA, USA). The cells were imaged by CLSM (Nikon, Tokyo, Japan) in two channels (ex/em: 488 nm/500-550 nm; 561/575-625 nm). The cellular uptake of sEV was quantified by measuring the RawIntDen (RID; a.u.) value in the red channel which is the sum of all fluorescent intensity in the region-of-interest (ROI) via ImageJ software.

### Combined ThS and Immunofluorescence Staining

0.6 mL of SH-SY5Y cells were plated at a density of 100,000 per well on the 12 mm poly-*L*-lysine coated round coverslips (Corning, NY, USA) in 24-well tissue culture plates and incubated overnight. After cell attachment, the coverslip was moved to another new well with 0.4 mL of medium on the second day. For seeding cells by mature tau fibrils, 10 µM of tau_P301L_ was incubated in an aggregation buffer for 4 days at 37 °C. Following incubation, 40 µL of tau fibrils was mixed with 160 µL of low-serum Opti-MEM medium and 10 µL of Lipofectamine 2000, and the mixture solution was incubated for 20 min at room temperature. Then, 50 µL of mixture solution was added to SH-SY5Y cells.

After 1 h adding tau fibrils, 50 μL of inhibitors were added to SH-SY5Y cells with *D*-GQD final concentrations around 0.3, 0.2, 0.1, and 0.05 µM in cells and incubated for 24 h at 37 °C. Inhibitors included *D*-GQDs-sEV, RVG-*D*-GQD-sEV, mxyl-NAP2/*D*-GQD-sEV, and RVG-mxyl-NAP2/*D*-GQD-sEV.

Briefly, The DOPE-PEG_3400_-RVG was incubated with the sEVs with a lipid: sEV ratio of 600:1 for 10 min at 37 °C. Then, the mixture solution was incubated with *D*-GQDs or mxyl-NAP2/*D*-GQD (8:1) complex (final concentration of *D*-GQDs and sEV: 8 µM and 1×10^9^ particles/mL) in 200 µL of PBS at room temperature for 20 min. sEVs were then washed with PBS (8 °C) three times under the support of a 100 kDa centrifuge tube to remove excess DOPE-PEG_3400_-RVG and unincorporated *D*-GQD or mxyl-NAP2/*D*-GQD complex.

After 24-h incubation, the cells were washed with PBS three times to remove tau fibrils and inhibitors, and fixed with 2% paraformaldehyde for 20 min. For ThS staining, fixed cells were incubated with 0.25% ThS for 8 min in the dark and washed three times with 50% ethanol. For immunofluorescence staining, the cells were incubated with 0.1% Triton X-100 for 15 min for permeabilization and washed three times with PBS. Following incubation in 5% BSA (1 h, room temperature) to block non-specific binding, the cells were covered with Tau46 (Santa Cruz Biotechnology, Inc., TX, USA; 1:700) diluted in 1.5% BSA overnight at 4 °C. The cells were washed three times for 5 min with PBS and then covered with m-lgG_1_BP-CFL 594 (Santa Cruz Biotechnology, Inc., TX, USA; 1:300) diluted in 1.5% BSA for 1 h at room temperature in the dark. Three PBS washes were performed for 5 min each and then the coverslips were mounted with mounting media. The cells were then imaged by CLSM (Nikon, Tokyo, Japan) in two channels (ex/em: 488 nm/ 500-550 nm; 561/575-625 nm). Tau fibrils in cells with green fluorescence signal from ThS were quantified with ImageJ software. The RawIntDen (RID; a.u.) value was measured in the green channel.

## Acknowledgements

We acknowledge funding from the National Institutes of Health (NIH 1R35GM15608-01 to Y.W. and NIH R01AG074570 to J.R.D.) and a STIR grant from the College of Engineering at the University of Notre Dame (to Y.W.). TEM and CLSM images were carried out in part in the Integrated Imaging Facility, University of Notre Dame, using Thermo Scientific Talos F200i and Nikon A1R-MP. Western blot images for immunodetection of proteins presented in this paper were acquired using the instrument of Biophysics Instrumentation Core/BIC, University of Notre Dame. We appreciate the support from these facilities for this study and would like to thank Maksym Zhukovskyi and Sara Cole for their knowledge and expertise as well as time towards this research. The authors also thank Hyunsu Jeon for the helpful discussion on the image analysis and are grateful to Kamlesh Makwana, Youwen Zhang, and James Johnston for their assistance with experiments. Fig. 1, 3e, and 5a were created using BioRender.com and used with permission.

## Data availability statement

The data supporting this article have been included as part of the supporting information.

