## Supplemental Information for "N-Amino Peptide-Graphene Quantum Dot Loaded Small Extracellular Vesicles for Targeted Therapy of Tauopathies"

### Supporting Figures

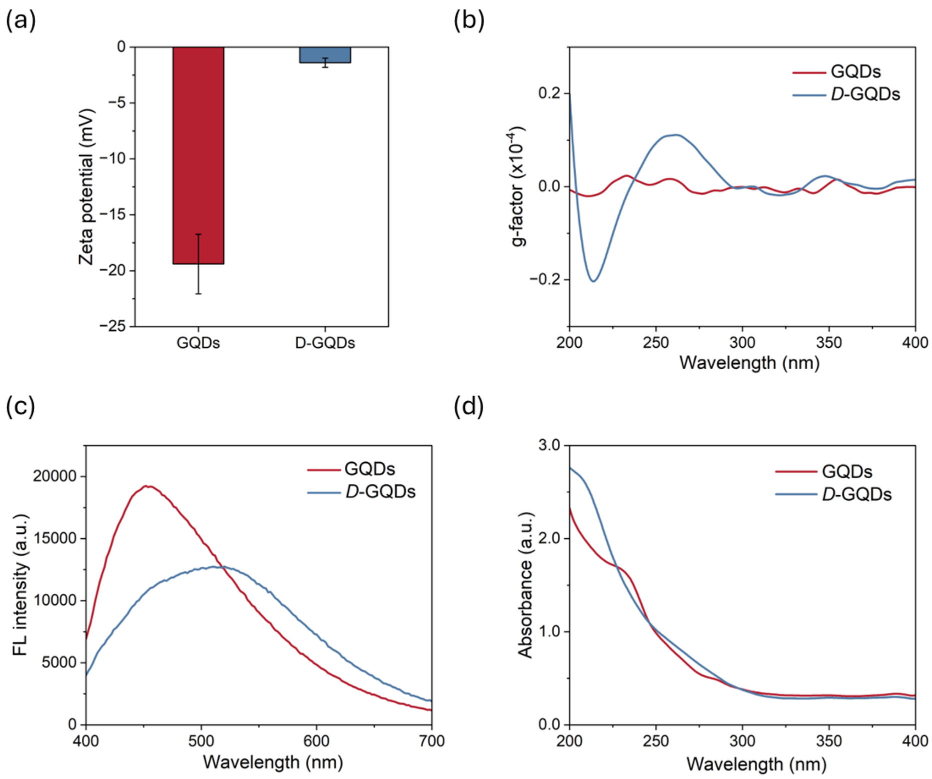


**Fig. S1** (a) Zeta potentials, (b) g-factor spectra, (c) fluorescence spectra (excited at 365 nm), and (d) absorbance spectra of graphene quantum dots (GQDs) and *D*-cysteine functionalized GQDs (*D*-GQDs).

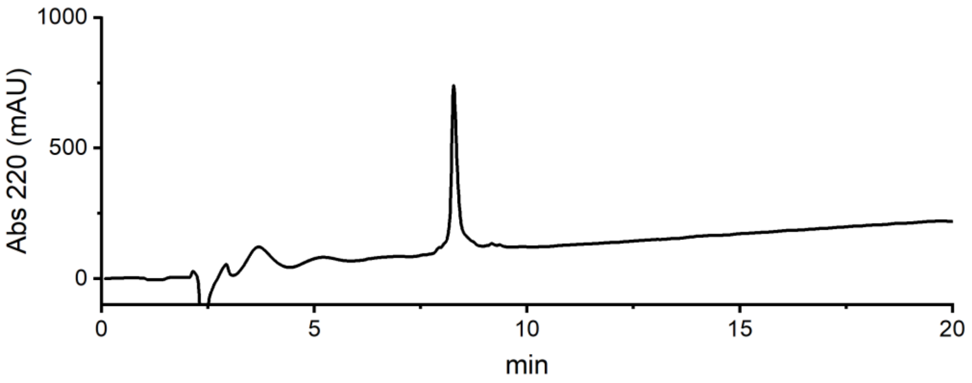


**Fig. S2** Mxyl-NAP2 purification and characterization. The crude peptide was purified by preparative scale RP-HPLC using a 5-80% MeCN/$H_2O$ gradient (with 0.1% formic acid). The pure peptide was obtained in 12% overall yield based on initial resin loading. HRMS (ESI-TOF) m/z $[M + H]^+$ calcd for $C_{112}H_{174}N_{33}O_{27}S_2$ 2477.2692, found 2477.2719, err -1.1 ppm.

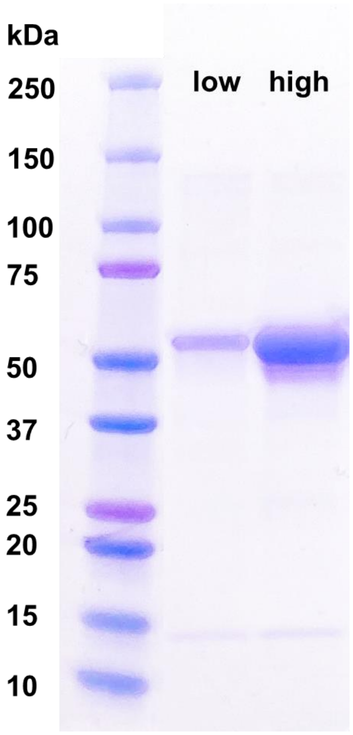


**Fig. S3** Sodium dodecyl sulfate–polyacrylamide gel electrophoresis (SDS/PAGE, Coomassie blue stain) of purified $tau_{P301L}$ protein loaded at low and high concentrations.

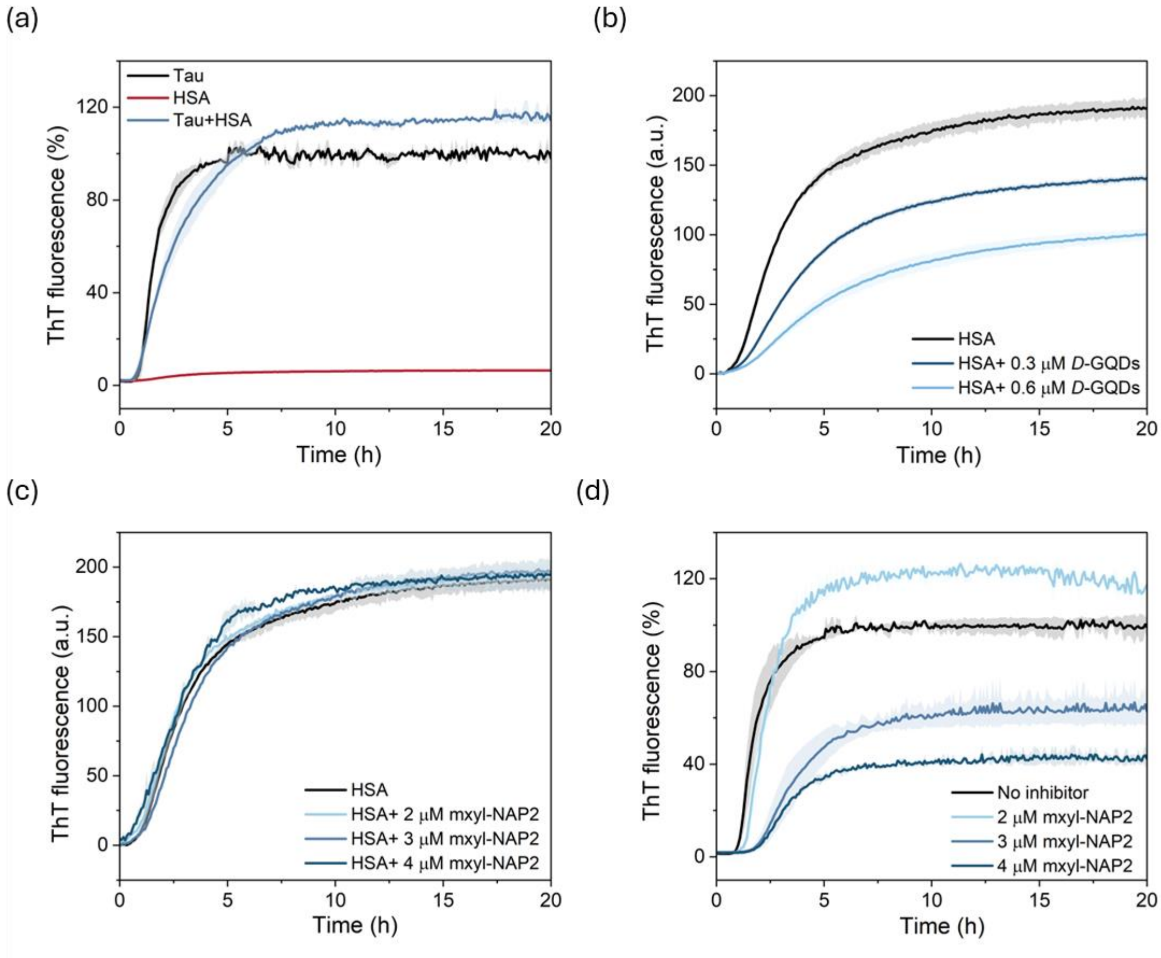


**Fig. S4** (a) Thioflavin T (ThT) fluorescence of $tau_{P301L}$ aggregation in the presence of human serum albumin (HSA), and $tau_{P301L}$ and HSA aggregation alone. (b-c) ThT fluorescence of HSA aggregation incubated with *D*-GQDs and mxyl-NAP2 at different concentrations. (d) ThT fluorescence of $tau_{P301L}$ aggregation incubated with different concentrations of mxyl-NAP2.

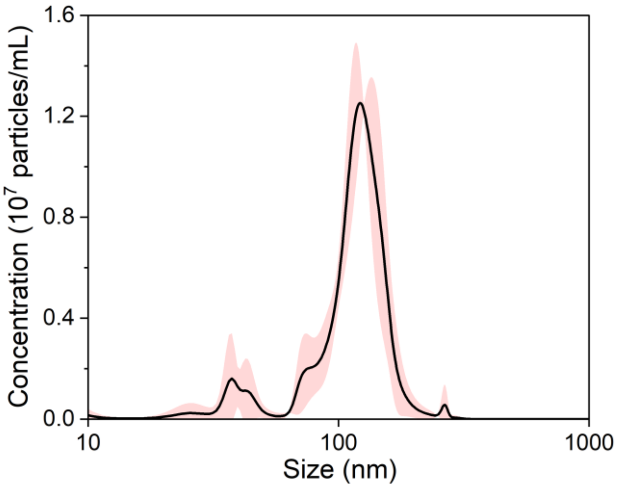


**Fig. S5** Nanoparticle tracking analysis (NTA) of *D*-GQD loaded small extracellular vesicles (sEVs) with an average diameter of 127.6 ± 3.1 nm.

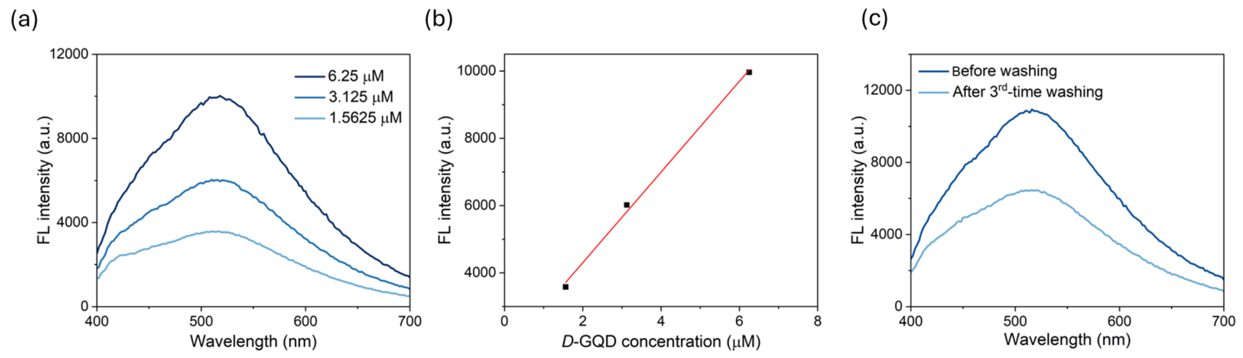


**Fig. S6** (a-b) The fluorescence spectra (excited at 365 nm) of *D*-GQDs at different concentrations. The fluorescence intensity at the emission peak exhibited a linear relationship with its concentration. (c) The fluorescence spectra (excited at 365 nm) of *D*-GQD loaded sEV before PBS washing and after three-time washing. Based on the concentration calibration curve of *D*-GQDs, *D*-GQD remained in the sEV solution after three-time PBS washing was 45% of the initial incubation concentration.

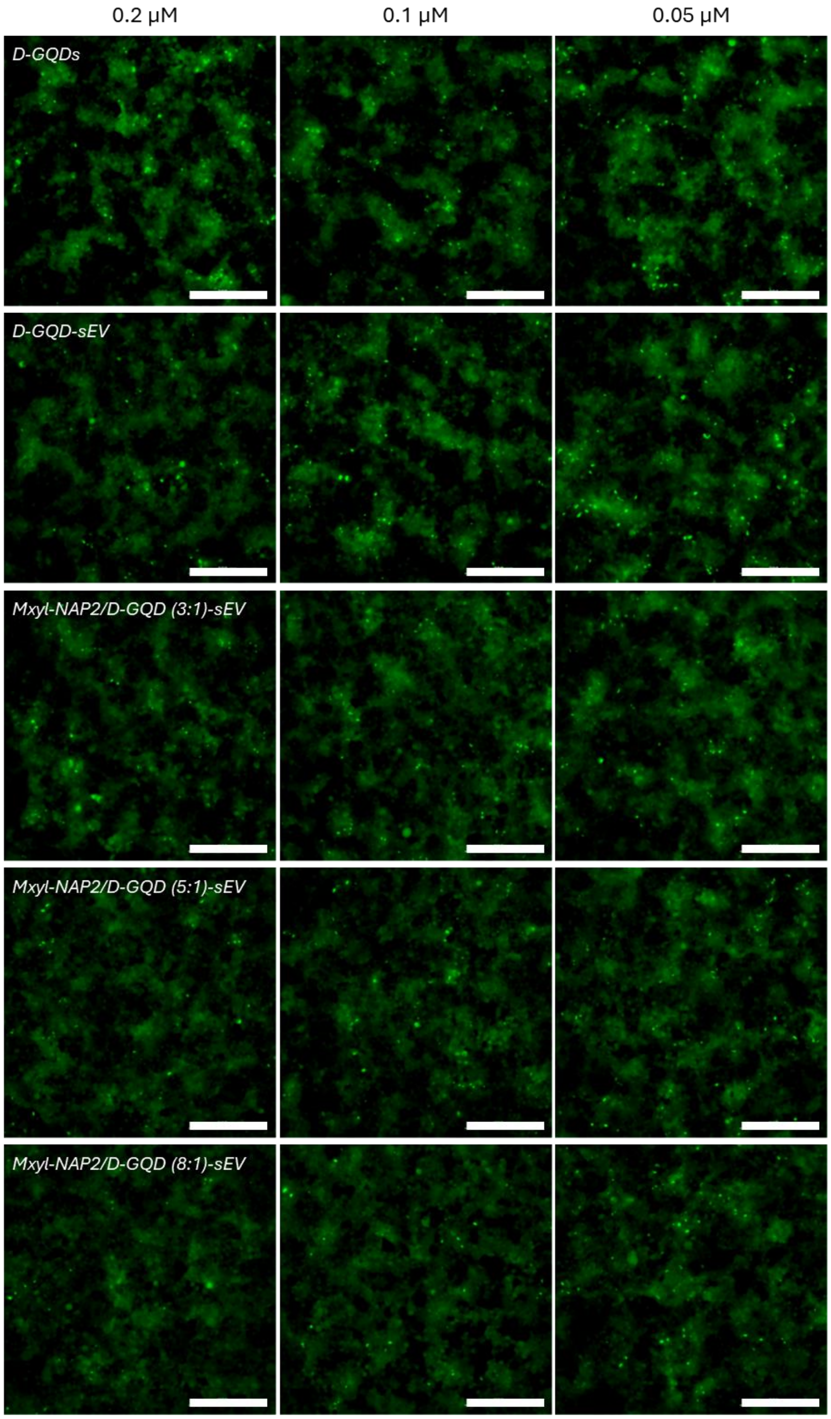


**Fig. S7** Mxyl-NAP2/*D*-GQD-sEVs, *D*-GQD-sEVs, and *D*-GQDs, at concentrations of 0.2, 0.1, and 0.05 μM for *D*-GQDs, sEV ($1\times10^9$ particles/mL), prevented the cellular transmission of mature $tau_{P301L}$ fibrils (0.19 μM). Representative images of cells were taken at 20× magnification under FITC channel (ex: 469 nm/em: 525 nm). The green puncta with high fluorescence represented the aggregation of tau in cells induced by exogenous tau fibers. Scale bar: 200 μm.

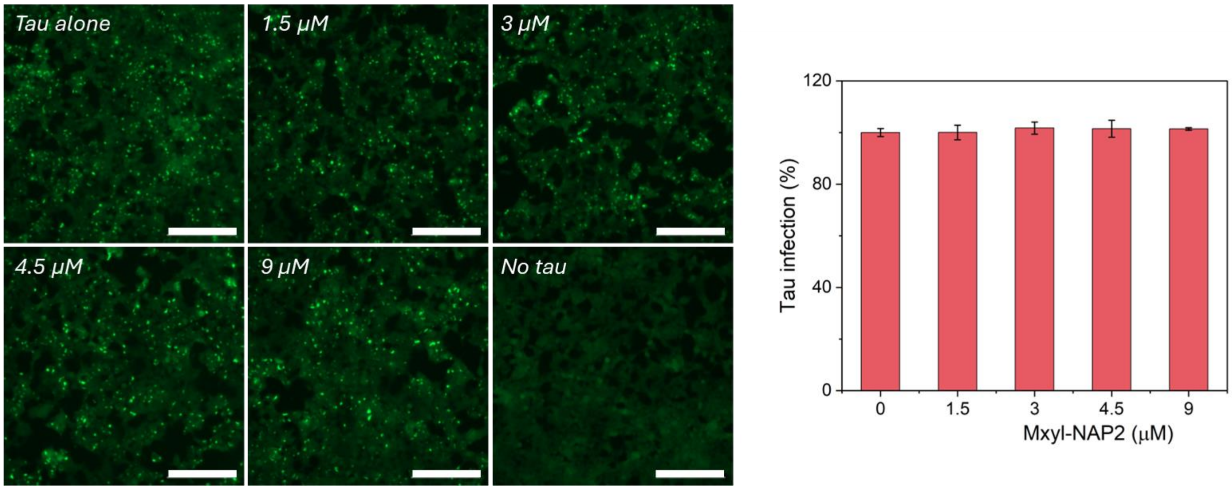


**Fig. S8** Mxyl-NAP2 was directly added to HEK293 cells stably expressing tau-RD (P301L/V337M)-YFP 1-h after adding $tau_{P301L}$ fibrils and incubated for 48 h. Mxyl-NAP2 at concentrations of 1.5, 3, 4.5, and 9 µM did not show the inhibtion ability on the cellular transmission of mature tau fibrils (0.19 µM). Fluorescent images of cellular tau biosensors were taken under the FITC channel (ex/em: 469/525 nm). The green puncta with high fluorescence represented the aggregation of tau in cells induced by exogenous $tau_{P301L}$ fibers. Scale bar: 200 µm. Tau infection (%) in the bar graph shows the number of intracellular fluorescent puncta relative to control infection wells lacking the inhibitors.

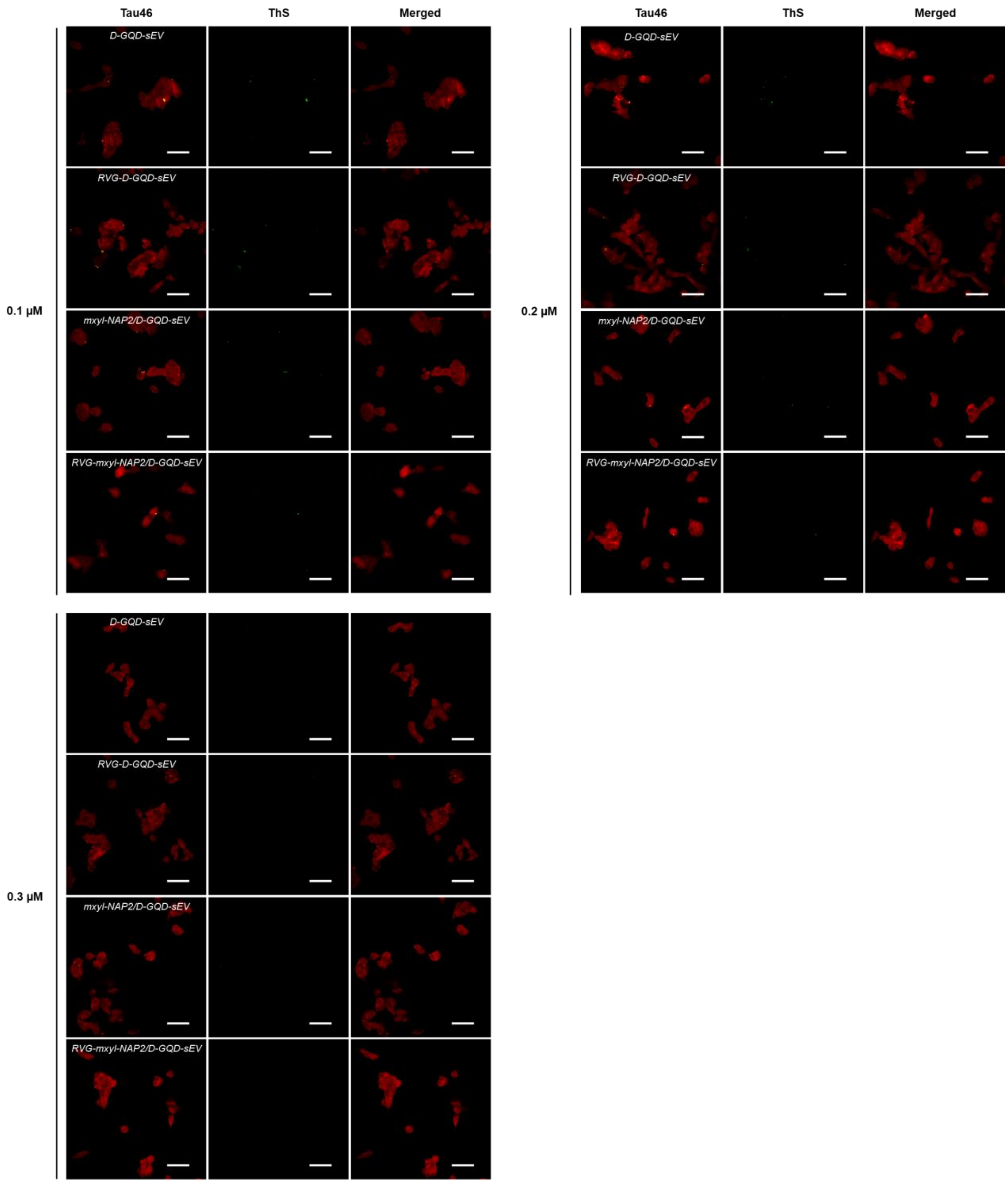


**Fig. S9** CLSM images of tau fibril-treated SH-SY5Y cells incubated with different inhibitors (*D*-GQD-sEV, RVG-*D*-GQD-sEV, mxyl-NAP2/*D*-GQD-sEV, and RVG-mxyl-NAP2/*D*-GQD-sEV) at concentrations of 0.1, 0.2, and 0.3 μM for *D*-GQDs. SH-SY5Y cells were sequentially stained with antibody Tau46 (red) for intracellular tau and Thioflavin S (ThS, green) for tau fibrils. Scale bar: 50 μm.

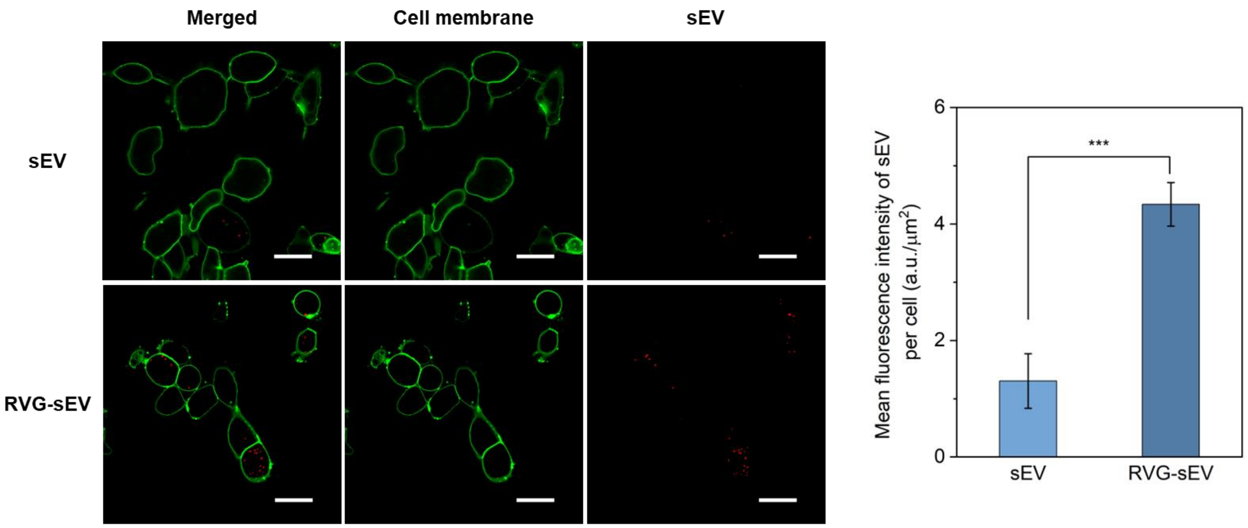


**Fig. S10** Cellular uptake profiles of sEV and RVG-sEV into SH-SY5Y cells. Each channel represents: green for CellBrite green cytoplasmic membrane dye and red for the DiI-stained sEV. SH-SY5Y cells were incubated with sEV and RVG-sEV (800 sEV/cell) for 4 h.

### Supporting Tables

**Table S1** Encapsulation efficiency of mxyl-NAP2 in sEV with different ratios of mxyl-NAP2 to *D*-GQDs was determined by the absorbance at 292 nm of encapsulated mxyl-NAP2 which was separated from *D*-GQDs and sEV membrane after lysis.

| | Abs at 292 nm | Concentration of mxyl-NAP2 | Encapsulation efficiency |
|---|---|---|---|
| Mxyl-NAP2/*D*-GQD (8:1)-sEV | 1.0336 | 24.67 μM | 30.8% |
| Mxyl-NAP2/*D*-GQD (5:1)-sEV | 1.0299 | 18.50 μM | 38.5% |
| Mxyl-NAP2/*D*-GQD (3:1)-sEV | 1.0249 | 10.08 μM | 42.0% |
